# Reevaluation of the *Toxoplasma gondii* and *Neospora caninum* genomes reveals misassembly, karyotype differences and chromosomal rearrangements

**DOI:** 10.1101/2020.05.22.111195

**Authors:** Luisa Berna, Pablo Marquez, Andrés Cabrera, Gonzalo Greif, María E. Francia, Carlos Robello

**Affiliations:** Laboratory of Host Pathogen Interactions-Molecular Biology Unit. Institut Pasteur de Montevideo. Montevideo, Uruguay; Laboratory of Apicomplexan Biology. Institut Pasteur de Montevideo. Montevideo, Uruguay; Departamento de Parasitología y Micología. Facultad de Medicina-Universidad de la República. Montevideo, Uruguay; Departamento de Bioquímica. Facultad de Medicina-Universidad de la República. Montevideo, Uruguay

## Abstract

*Neospora caninum* primarily infects cattle causing abortions with an estimated impact of a billion dollars on worldwide economy, annually. However, the study of its biology has been unheeded by the established paradigm that it is virtually identical to its close relative, the widely studied human pathogen, *Toxoplasma gondii*. By revisiting the genome sequence, assembly and annotation using third generation sequencing technologies, here we show that the *N. caninum* genome was originally incorrectly assembled under the presumption of synteny with *T. gondii*. We show that major chromosomal rearrangements have occurred between these species. Importantly, we show that chromosomes originally annotated as ChrVIIb and VIII are indeed fused, reducing the karyotype of both *N. caninum* and *T. gondii* to 13 chromosomes. We reannotate the *N. caninum* genome, revealing over 500 new genes. We sequence and annotate the non-photosynthetic plastid and mitochondrial genomes, and show that while apicoplast genomes are virtually identical, high levels of gene fragmentation and reshuffling exists between species and strains. Our results correct assembly artifacts that are currently widely distributed in the genome database of *N. caninum* and *T. gondii*, but more importantly, highlight the mitochondria as a previously oversighted source of variability and pave the way for a change in the paradigm of synteny, encouraging rethinking the genome as basis of the comparative unique biology of these pathogens.

## INTRODUCTION

The Apicomplexa comprise a large phylum of parasitic alveolates of medical and veterinary importance, causing deadly diseases such as malaria, criptosporidiosis, neosporosis and toxoplasmosis, among others. With the exception of a few commonalities such as their obligatory intracellular lifestyle, the presence of specialized secretory organelles and of secondary endosymbionts, the apicomplexans differ greatly in morphology, host range specificity, pathogenicity, reproductive strategy and transmission. Understanding the molecular basis of these differences has been the focus of much research. Comparative genomic analyses revealed that, albeit all small, apicomplexans genomes vary greatly in size, ranging from 9 to 130 Mb^1,2^. Having diverged from a common ancestor 350-824 million years ago^3^, shy of 900 genes are conserved amongst them, whereby major genomic rearrangements can be observed^1^.

High synteny, defined as conserved content and order of a given genomic locus, is rarely observed^1^. A seemingly stark exception to this are the genomes of *Toxoplasma gondii* and *Neospora caninum.* Morphologically, these parasites are virtually indistinguishable, so much so, that *N. caninum* was only recognized as a separate species in 1988^4,5^. Moreover, both species exhibit similar tropism within their hosts, where they can infect virtually any nucleated cell. They both exhibit a fast replicating form (tachyzoite) causing acute disease, that transitions into a slow dividing form (bradyzoite) which persists in immune-privileged sites, such as the brain, establishing chronic infection. In line with this, initial comparative analysis concluded that these species have largely conserved genomic content, and are largely syntenic^6^. Despite their commonalities, however, the biology of these pathogens also differs significantly. *T. gondii* infects a wide range of intermediate hosts, including humans, causing deadly disease in immunocompromised individuals or by congenital transmission. In contrast, *N. caninum* infects primarily cattle, causing abortions with an estimated impact of a billion dollars on worldwide economy, annually^7^. Feline species act as definitive hosts of *T. gondii* whilst sexual replication of *N. caninum* occurs only in canids ^8–10^. These biological differences have been largely ascribed to absence, point mutations and pseudogenization of *T. gondii* virulence factors in *N. caninum*, and the comparative amplification of surface protein-coding gene families in *N. caninum*^6,11^.

Advancements in genome sequencing technologies have accompanied the fast-paced genomics era. Particularly, third generation sequencing technologies, such as Pacific Bioscience (PacBio) and Oxford Nanopore sequencing outperform prior technologies by providing very long reads that can span regions containing repetitive sequences. This has led to improvements in the assembly of previously unattainable genomes, such as those presenting high proportions of repetitive sequences, allowing whole new genomes to be assembled with high accuracy. Here, we set out to sequence and *de novo* assemble two *N. caninum* strain’s genomes and the *T. gondii* genome, using PacBio and Oxford Nanopore. Our *de novo* assembly of the three genomes identifies largely incorrectly assembled reference genomes. Strikingly, our work reveals that previously annotated chromosomes VIIb and VIII are in fact a single chromosome in both species, reducing the total number of chromosomes in these coccidia to 13. Greater sequencing coverage and corrected assembly uncovers major chromosomal rearrangements, hundreds of previously unidentified *N. caninum* genes, and more accurate account for gene copy number. In addition, we fully annotate the previously uncharacterized apicoplast genome of *N. caninum*, and note large levels of mitochondrial heteroplasmy exist in both species. More importantly, our results ultimately challenge the paradigm of gene content synteny between *T. gondii* and *N. caninum*, prompting us to further explore these species unique genome structures as potentially overlooked sources of important biological information.

## MATERIAL AND METHODS

### Cell culture

*N.c. Liverpool* was acquired from ATCC (50845). *N.c.Uru1* was isolated from a local congenitally infected calf as described in (^12^). *T. gondii* RH ΔKu80^13^ strain was kindly provided by Boris Striepen. All strains were maintained and grown as described in Cabrera, et. al. 2019^12^.

### High Molecular Weight DNA extraction and sequencing

For “Oxford Nanopore” and Illumina sequencing, DNA was extracted from *N. caninum* Liverpool, *N. caninum* Uru1 and *T. gondii* RH ΔKu80^13^ strain, using a DNA purification kit from Zymo Research (#D4074). Minion Oxford Nanopore sequencing was done *In house* as described in ^14^. Sequencing libraries were prepared using a ligation sequencing kit and a Native barcoding expansion kit (EXP-NBD103/SQK-LSK108, Nanopore, England) according to (^15^), starting from 1 μg of total genomic DNA. 12 hours sequencing was performed in an R9.4 Flow Cell (FLO-MIN106, Oxford Nanopore). Base calling and sequence retrieval was done using Guppy basecaller version 3.0.3. For PacBio sequencing, DNA was extracted from *N. caninum Liverpool*, *N. caninum Uru1* and *T. gondii* RH ΔKu80^13^ strain by overnight incubation in lysis buffer (Tris-EDTA buffer supplemented with 1 ug/mL of RNAse A, 10% SDS, 20% Proteinase K) at 55°C, phenol/chloroform extraction, ethanol precipitation and resuspension in ultrapure water. PacBio Sequencing was performed at the Integrative Genomics Core (Beckman Research Institute, Monrovia, CA.), using 5 SMRT cells per sample for *N. caninum* and *T. gondii* genomes. Illumina sequencing of *N. caninum* DNA was also done at the Integrative Genomics Core to 65x coverage.

### Genome assembly and annotation

PacBio reads were assembled using HGAP Assembly software^16^. Oxford Nanopore reads were assembled using CANU^17,18^. Assemblies were merged using Quickmerge software^19^. A single conflict region in the assembly was solved by PCR using primers: M1_F: GAGGCGCTTACAATCAACCC, H37_F_M1_R: GAGACAGGACGGACTGAAGA, H37_R: CTGCTCTGTCTGAACAGGTT, M37_F: GCGAACAGCACGAAGTGAGA, M37_R: TCGTGCTTTGAGCATCCTCT. Short nsertions and deletions, a common artifact produced by long read technologies, were corrected using illumina reads in Pilon^20^. Reads used include our own, from DNA purified as described above, as well as raw reads obtained from Sequence Read Archive (SRA) NCBI repositories (Accession IDs: PRJNA531306, ERR012899 and ERR012900). Gene annotation was performed using the automated annotation tool COMPANION^21^ using Agustus Threshold 0.2, Taxon ID 5811, Align reference proteins to target sequence as parameters (others as default), and supporting RNA seq data to produce a transcript assembly. This was first aligned with Cufflinks^22^ and assembled with TopHat^23^ using SRAs with accession IDs ERR690607, ERR690608, SRR4013168, SRR4013169, SRR4013170, SRR4013171, SRR4013172 and SRR4013173 as input sequences. Apicoplast and mitochondrial genomes were identified by manual GC filtering and confirmed by Blast. Apicoplast genome was annotated using Mfannot^24^. Mitochondrial genome annotation was done in MITOS (http://mitos2.bioinf.uni-leipzig.de/index.py) using Opisthokont as reference. PacBio and MinION data has been deposited in NCBI repository (BioProject ID PRJNA597814).

### Comparative Genomic Analysis

Assembled genomes were compared using NUCmer^25^ to create the alignments between the assemblies being compared and Assemblytics^26^ for visualization. Plots comparing the synteny between assemblies were obtained with the visualization tool Circos^27^, using the output from Blast to create links between chromosomes. Repetitive regions were analyzed using YASS^28^. Individual chromosome comparisons were performed with Artemis^29^ and ACT. IGV^30^ was used for visual inspection of aligned reads (wgs and rna-seq) on the assemblies. Specific scripts generated in this study were written in R environment^31^ and Bash to parse results and automate pipelines. Telomeres were identified by searching on chromosome ends for the typical TTTAGGG and AAACCCT septameric repeats. Centromeric regions in *Neospora* were determined by BLAST against previously identified centromere sequences in *Toxoplasma gondii* by ChIP-chip^32^

## RESULTS

### *Neospora caninum* and *Toxoplasma gondii* long-read assembly genomes

To assemble the *Neospora caninum* genome, we sequenced DNA from the reference strain *N. caninum* Liverpool (NcLiv), and a recent isolate from an experimental farm in Uruguay, named *N. caninum* Uru1 (NcUru1)^12^. Sequencing was done by Pacific Biosciences technology, Oxford Nanopore and Illumina. Reads derived from each sequencing were assembled independently, and then combined into a single assembly per strain (**SFig 1A**). For *NcLiv*, PacBio and Nanopore assemblies matched completely, with the exception of a single conflict region, which was resolved by PCR (**SFig1B-C**). The assembled genomes were corrected with publicly available Illumina reads or those obtained *in house*. For *NcLiv*, the sequencing resulted in over 100x depth, vastly improving the currently available genome coverage (**Table 1**). Genomes of *N. caninum* strains were assembled separately.

**Figure 1.**
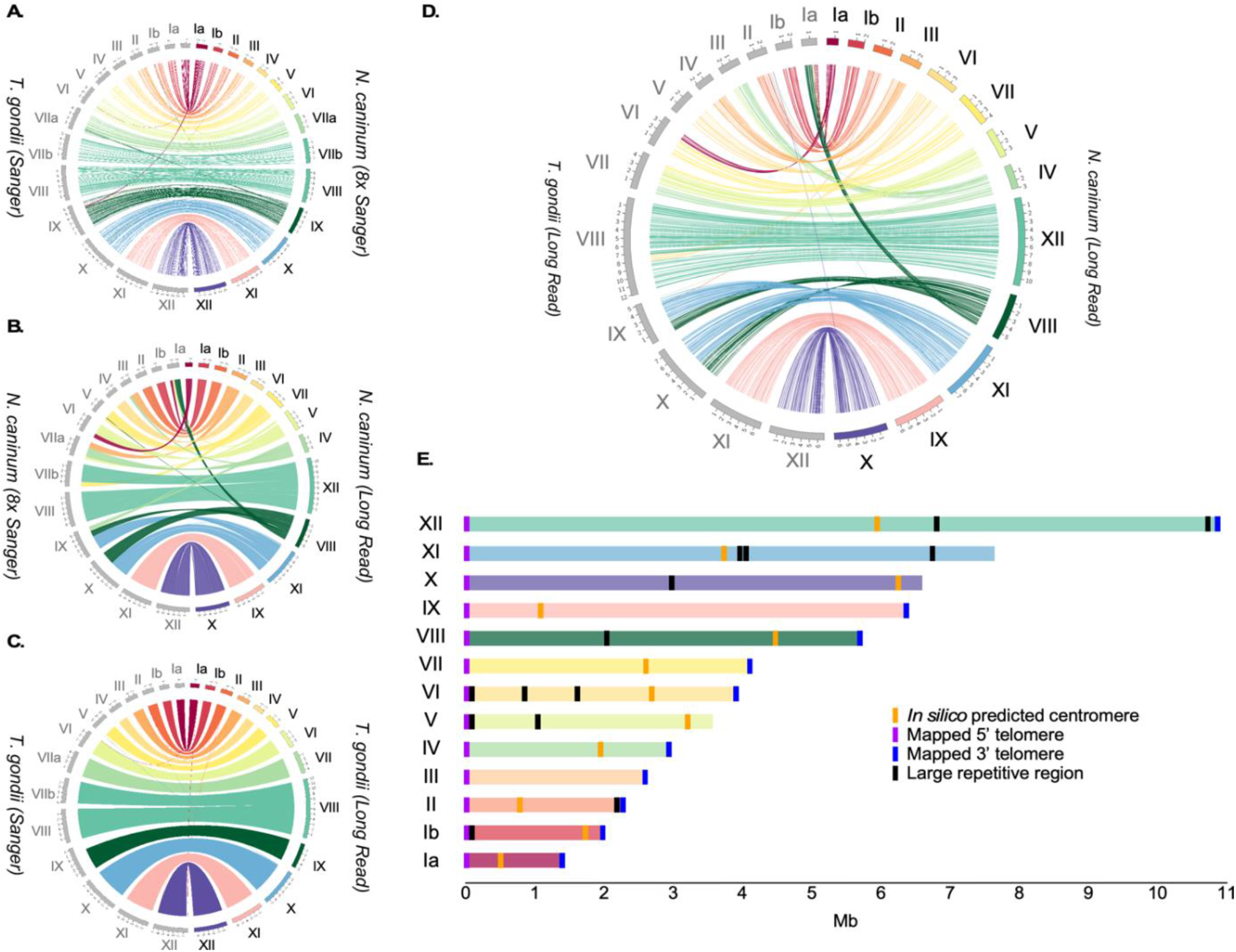
Comparative analysis of genome assemblies of *Neospora caninum* and *Toxoplasma gondii* using third generation sequencing data reveals misassembly and karyotype differences. **A**. Comparative analysis of the *Toxoplasma gondii* type II (TgME49) genome assembly and the *Neospora caninum Liverpool* strain genome assembly, obtained based on Sanger technology sequencing data. **B**. Comparative alignment of the *Neospora caninum Liverpool* genome assemblies using Sanger and third generation (long read) technology. **C**. Comparative alignment of the *Toxoplasma gondii* type II (TgME49) genome assemblies based on Sanger technology sequencing data or third generation (long read) technology of *Toxoplasma gondii* type I (TgRH). **D.** Comparative alignment of the *Toxoplasma gondii* type I (TgRH) and the *Neospora caninum Liverpool* genome assemblies based on third generation (long read) sequencing technology. **E.** Chromosomal layout of Neospora caninum. Karyotype, chromosome length, telomeres, putative centromeres and large repeats, are shown.

**Table 1.** Metrics of the *de novo* genome assembly of *N. caninum* Liverpool.

| Nuclear Genome (NcLiv) |  |  |
| --- | --- | --- |
| Genome properties |  |  |
|  | Chromosomes | 13 |
|  | Unplaced contigs | 31 |
|  | Total length | 61.5Mb |
|  | GC (%) | 54.8 |
|  | N50 | 6.4Mb |
|  | N75 | 3.6Mb |
|  | L50 | 4 |
|  | L75 | 8 |
|  | # N's per 100 kbp | 0 |
| Protein-coding genes |  |  |
|  | Number of gene models | 7540 |
|  | Pseudogene | 258 |
|  | Mean exons by gene | 5.8 |
|  | Percentage coding | 59.2% |
|  | Mean CDS length (pb) | 4872 |
|  | Mean exonic length (pb) | 423 |
|  | Coding GC (%) | 60.1% |
| **Intergenic regions** |  |  |
|  | Mean Length (pb) | 3091 |
| **RNA genes** |  |  |
|  | tRNA | 157 |
|  | rRNA genes | 22 |
|  | snRNA | 12 |
| **Apicoplast Genome (NcLiv)** |  |  |
|  | Contigs | 2 |
|  | Total length | 52.3 Kb |
|  | GC (%) | 23.7% |
| **Mitochondrial Genome (NcLiv)** |  |  |
|  | Contigs | 9 |
|  | Largest contig | 32.4 Kb |
|  | Total contigs length | 121 Kb |
|  | GC (%) | 36.8% |

Assemblies for both *Neospora* strains were practically indistinguishable, indicating high genome similarity. *NcLiv* genome consisted of 13 large contigs and 31 short (less than 122kb) unplaced fragments. With an N50 of 6.4Mb, 75% of the 61.6 Mb nuclear genome is in 8 of the largest chromosomes **(Table 1).** The 13 largest contigs correspond to complete chromosomes; both 5’ and 3’ telomeres were mapped for 10, while we found one telomere at one end and sub-telomeric regions at the other end for 3 (**Supp.Table 1**). Putative centromeric sequences were identified *in silico* either by blasting the *T. gondii* centromeres (Chr Ia, II, IV and V) or by identifying large regions devoid of gene-coding sequences flanked by syntenic genes flanking the centromeric regions of *T. gondii (*Chr Ib, IX and XII) (**Fig 1B-E** **and Supp. Table 2**). Our assembly revealed that, in contrast to what has been reported, the *N. caninum* genome is not organized in 14 chromosomes. Rather, previous ChrVIIb and ChrVIII are in fact a single chromosome (**Fig1 B)**.

Given that the previous *N. caninum* genome assembly had been constructed using as the reference the *Toxoplasma gondii* genome, by assuming that these species conserve synteny and gene order, we next examined how our new *N. caninum* genome assembly compared to that of the *T. gondii* reference genome**(Fig 1A** ). To our surprise, not only did the newly assembled *N. caninum* genome differ from that of *T. gondii* in the total number of chromosomes, but a number of large chromosomal rearrangements could be observed (**Fig1 B)**. Strikingly, only half of the chromosomes’ structure, corresponding to *N. caninum*’s chromosomes Ib, II, III, IX, X and XII, are maintained between these species.

Next, we wondered whether the observable differences between the species were an artifact of the *T. gondii* genome assembly based on shorter reads. To this end, we sequenced the genome of a Type I strain (RH) of *T. gondii* both by PacBio and Oxford Nanopore, and *de novo* assembled the genome. Our long read sequencing resulted in a 108x coverage, and its assembly closely matched that reported by using Sanger and 454 sequencing (**Fig1 C)** confirming that the differences observed with *N. caninum* are not artifactual. However, we were able to once again observe the fusion of chromosomes VIIb and VIII in *Toxoplasma*, thereby indicating that the karyotype for both these apicomplexans is 13, and not 14 as previously reported (**Fig1 D)**.

We next explored whether the breaks in synteny between these closely related species correlated with any distinguishable genomic features. We noticed that the level of sequence identity between the species varies along chromosomes. Coding regions seem to be highly conserved (82.4%). Nonetheless, we noticed that there is a detectable shift in codon usage between the two species, whereby *N. caninum* tends to use GC richer codons than *T. gondii.* The difference in codon usage conveys a detectable difference in composition whereby the mean GC content for N. caninum is 58.3% while it is 56.4% for *T. gondii*, being this a statistically significant difference (*p-value* 1.4×10^−9^). The same trend can be observed for GC3 (63.7% and 61.0%, respectively) and less markedly for GC1 (**SFig 2**). On the other hand, non-coding regions, including intronic and intergenic regions are on average 80.7% identical. Non-coding regions throughout the genome of *N. caninum* are interspaced with low complexity repetitive sequences rich in A/G or A/T. In addition, we identified three conserved sequence motifs present at a number of regions where synteny is interrupted. Particularly, five chromosomes (IV-VIII) exhibit comparative chromosomal rearrangements at regions that coincide with repetitive units of such domains (**Fig2A** **and** **B**, and **Supp. Table 3**).

**Figure 2.**
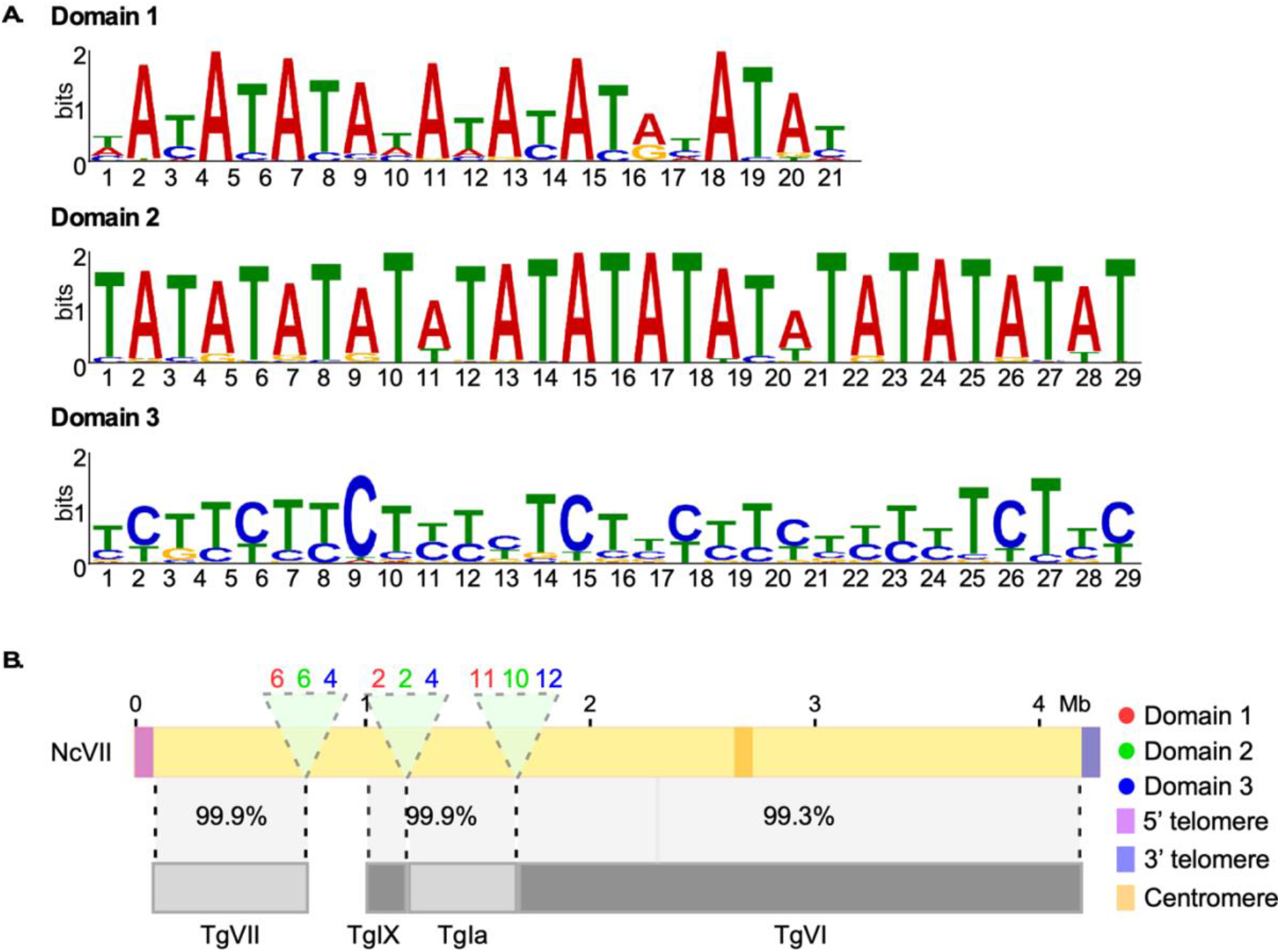
Regions of synteny breaks between *N. caninum* and *T. gondii* are populated by three conserved domains. **A.** Sequence identity of domains identified at regions where chromosomal rearrangements have occurred. **B. Graphical representation of chromosome VII of** *N. caninum Liverpool*. Comparative alignment to the *Toxoplasma gondii* chromosomes. Percentages of sequence identity are shown. Regions examined for the presence of motifs are indicated (a, b, and c, light green). The position of the putative centromere is indicated in orange. Note that large repetitive regions were not identified in this chromosome. 5’ (light purple) and 3’ (dark purple) telomeres are indicated. Identity and number of domains found per region, in chromosome VII, are indicated.

### Gene Annotation

Annotation of the newly assembled NcLiv genome, using homology-based searches and RNA-seq datasets, resulted in the annotation of 7540 genes and 254 pseudogenes. Of these, 7502 are distributed in 13 chromosomes while 38 are in unplaced sequences. The products of 4553 genes have a putative assigned function, while 2987 code for hypothetical proteins. Comparatively, 554 new genes were identified in our annotation; 9 are coded for in unassembled sequences, while 545 are distributed in the chromosomes. Albeit not originally annotated, 516 of these can be found in the original Sanger sequence reads of the *NcLiv* genome, with over 95% identity in over 95% of the gene’s length. Of these, 494 mapped to the originally assembled chromosomes, while 22 belong to unplaced sequences.

No common theme was observed among the newly annotated genes. Of these, the most frequently found are SRS domain-containing proteins, followed by WD-domain, and zinc-finger containing proteins. This is consistent with previous findings showing that the SAG-related family of surface proteins is amplified in *N. caninum*^6^. Previously, 227 members of this family had been identified. Here, we annotate a total of 231 SAG-related proteins.

Finally, 38 novel genes were uniquely identified in the newly sequenced genome; 36 are distributed in 11 of the assembled chromosomes, while 2 remain in unplaced sequences (**Fig3 A**). Of these, 6 have putative assigned functions or recognizable domains; a Syntaxin 6, N-terminal/SNARE domain containing protein, an SPX domain/VTC domain containing protein, a Kelch motif/Galactose oxidase, central domain/BTB/POZ domain containing protein, a glutathione S-transferase, N-terminal domain containing protein, and a Cyclophilin type peptidyl-prolyl cis-trans isomerase (**Fig3B** **and Supp. Table 4**). The products of the remaining 32 genes are regarded as hypothetical. We note that many of this new genes resulted either from the annotation of sequencing gaps (**Fig 3A**) or sequencing that extended through repetitive regions where previous sequencing had failed (See **Fig3C** for a representative example)

**Figure 3.**
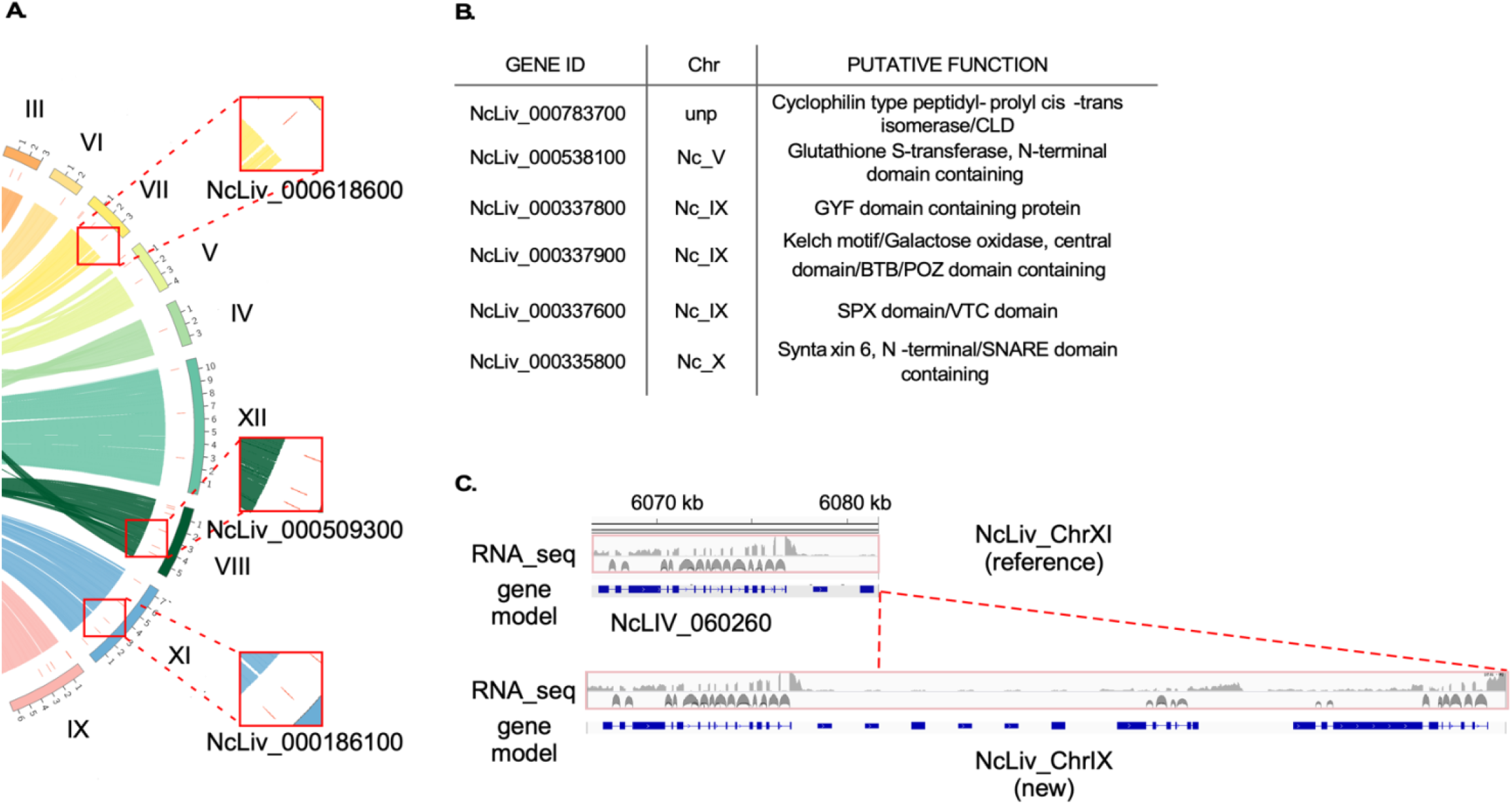
Gene annotation reveals previously unknown genes in the genome of *N. caninum*. **A.** Graphical representation of the position of novel genes in the newly assembled chromosomes. Red lines mark the position of novel genes along chromosomes. Alignment to previously assembled NcLiv genome is partially shown for reference. Three insets are shown to highlight the annotation of three new genes in three newly assembled genomic regions. **B.** Putative function of xix out of the 38 newly identified genes. The remaining 32 genes are annotated as hypothetical**. C.** Representative example of the improvement in annotation in regions that had been previously collapsed due to the presence of tandem repeats. Several new genes were annotated, all of whose annotation is supported by RNA-seq data.

Next, we explored whether the new annotation revealed presence of uncharacterized virulence factors. We surveyed the genome for homologs of major virulence factors characterized in *T. gondii*, the majority being kinases involved in protecting the parasitophorous vacuole from host-cell intrinsic defenses; ROP5, ROP16, ROP17, ROP18 and ROP38, dense granule secreted effectors such as GRA16, GRA24, GRA25 and GRA44, as well as, IST. No major differences were found between the annotations regarding virulence factors with the exception of ROP38, whereby 4 gene copies had been reported, we could resolve 9 ROP38 copies arranged in tandem. Our annotation is consistent with previous reports describing the absence of GRA24 and IST homologs in *N. caninum*.

### *Neospora caninum* apicoplast genome

The apicoplast is a relic non-photosynthetic chloroplast-like organelle of bacterial origin present in most apicomplexan parasites, whereby essential lipid synthesis occurs^33,34^. This organelle is of great importance as it has been validated as the target of anti-parasitic drugs such as clindamycin^34^. Most apicoplast proteins are encoded for in the nucleus, and later imported^35–41^. However, the apicoplast harbors its own genome, which has been traditionally regarded as coding for proteins needed for its maintenance^42^.

Surprisingly, we identified two contigs of 44 and 26kb, with markedly lower GC content than that of the nuclear genome (21.5 vs. 59.2%), and a 10kb inverted repeat sequence (**Fig 4A-H**). Annotation of these revealed 60 open reading frames most of which correspond RNA polymerase subunits and ribosomal proteins. In addition, the coding regions for SulfB and Clp protease, and a hypothetical protein were identified (**Fig4C, E** and **Supp. Table 5**). These have been shown to be resident proteins of the apicoplast, thereby indicating these two contigs correspond to the apicoplast genome of *N. caninum*.

**Figure 4.**
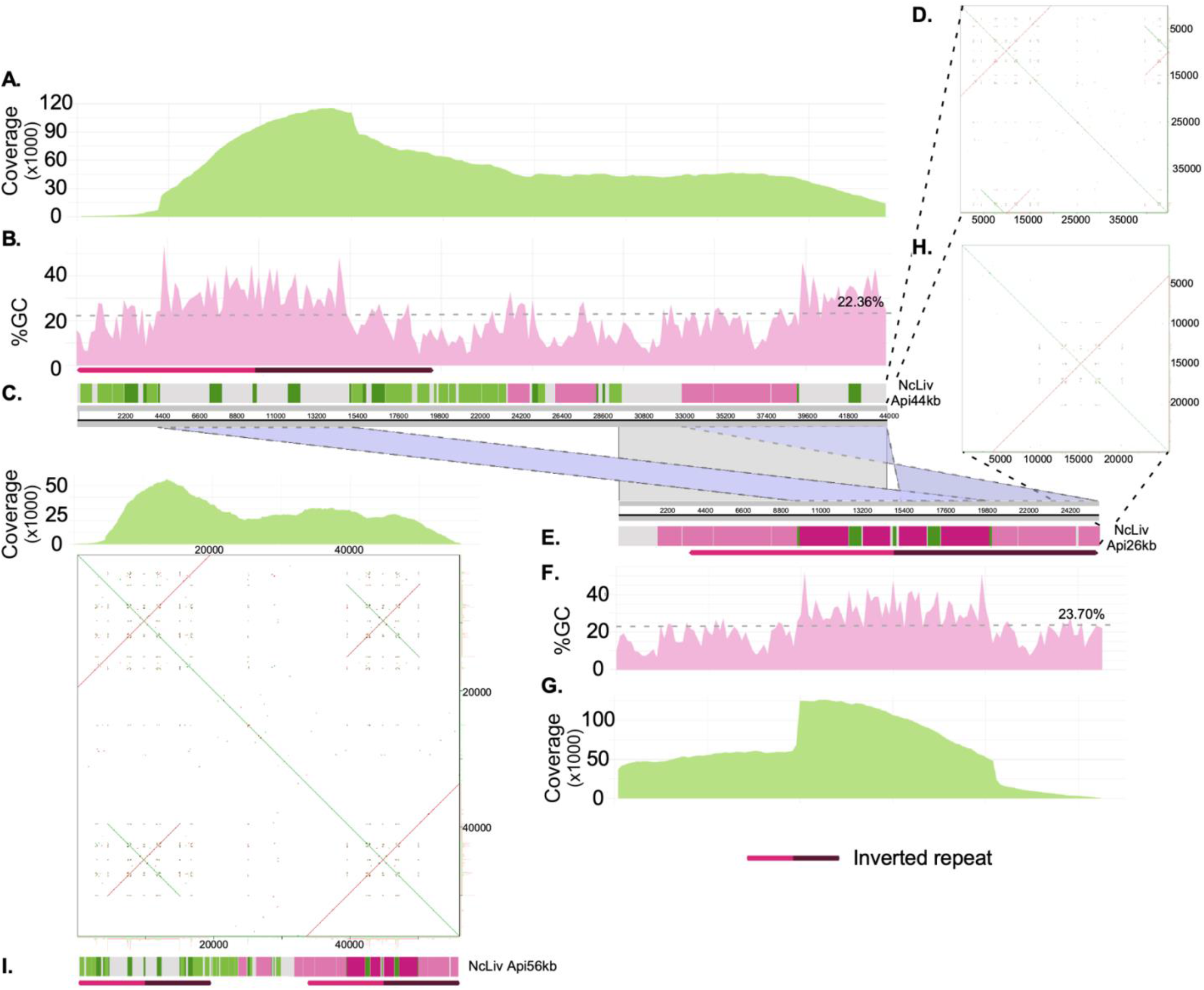
*Neospora caninum* Apicoplast genome structure and annotation. **A.** Read Coverage count along the length of the 44kb apicoplast genome contig. **B.** Percentage of GC along the length of the 44kb apicoplast genome contig. Average GC is shown above the dotted line. **C.** Annotation along the length of the 44kb apicoplast genome contig (see supplementary Table 5 for details) **D.** The presence of an inverted repeat sequence at the 5’ end of the contig is graphically represented in a YASS plot. **E.** Annotation along the length of the 26kb apicoplast genome contig (see supplementary Table 5 for details) **F.** Percentage of GC along the length of the 26kb apicoplast genome contig. Average GC is shown above the dotted line. **G.** Read Coverage count along the length of the 26kb apicoplast genome contig **H.** The presence of an inverted repeat sequence at the 3’ end of the contig is graphically represented in a YASS plot. **I.** Graphical representation of the apicoplast genome structure if both contigs are fused together Read Coverage count supporting the fusion, along the length of the resulting 56kb apicoplast genome contig, are shown. Note that both annotation and the position of the inverted repeat sequences are shown. The YASS plot graphically shows the presence of both the 5’ and 3’ inverted repeats. Note, however, that the presence of this highly similar repetitive elements at both ends is not sufficient to support circularization of the genome.

The apicoplast genome has been reported to be circular. However, we do not find compelling evidence to favor circularization of the apicoplast genome over a two linear chromosome model, in spite of using long reads, regarded as optimal to resolve these kinds of conundrums (**Fig 4I** **and SFig 3**). The assembled sequences bear significant homology at their ends, allowing their collapse into a single molecule of 56kb. However, mapped reads to this pseudomolecule do not support its circularization (**Fig4I** **and SFig 3C)**. This could be explained by lower sequencing coverage of the apicoplast genome due to its markedly lower GC content, or alternatively, it could suggest that the genome is indeed linear.

### Mitochondrial genomes

A number of large contigs (up to 32.4kb) amounting to a total of 121kb were identified upon sequencing *NcLiv* (**Table 1**, **Fig5** **and SFig 4**). These contain sequences with homology *to* classically known mitochondrial genes such as cox1 and cob1, and ribosomal RNAs (**SFig 4 and Supp Table 6**). In addition, these contigs display a GC content averaging 36.8%, markedly lower than that of the nuclear and apicoplast genomes (**Table 1**).

Interestingly, we could not assemble these contigs together, and neither one of them circularizes on its own. All of these contigs contain fragments of cox1 and cob1 open reading frames, however, the vast majority of them are interrupted by internal STOP codons. Three of the contigs feature replication origins of the heavy strand (OH) sequences, and only one of them features a potentially functional cox1 gene with no internal STOPs. One other contig lacks an origin of replication but features a potentially functional cob1 copy (**Supp Table 6**).

The mitochondrial genome of *NcUru1* is distributed in 20 contigs, which vary in size ranging from 1437bp to 86kb with a medium of 12kb in size, and average a GC contents of 36.95%. The largest contig, of 86 kb, features three origin of replication (OH sequences), 36 fragments with homology to cytochrome b, 56 fragments with homology to cytochrome c oxidase subunits I and III, and several ribosomal RNA encoding genes. All genes encoding for cob, cox1 and cox3 contain internal STOP sequences which would result in truncated proteins. Likewise, both genomes encode for the intronic endonuclease LAGLI.

Interestingly, while apicoplast genomes are 95% identical between the two *N. caninum* strains sequenced, the contigs corresponding to the mitochondrial genomes of *NcUru1* and *NcLiv* did not coincide with each other (**Fig5**). They both showcase a similar structure consisting of multiple linear contigs, featuring pseudogenized copies of cox1 and cob1, whose order seems to reshuffle in every contig (**Fig5**). The gene order, distribution and length is unique to each strain.

Twenty nine contigs ranging from 1.1 to 39 kb in size, with a median of 8.0kb were identified as the mitochondrial genome of *T. gondii*. The contigs’ GC content averages 36.6%. A similar structure of shuffled pseudogenes can be observed, whereby multiple copies of different size coding fragments of cob, cox1 and cox3 populate the mitochondrial chromosomes, together with rRNAs (**Fig5**). Origins of replication of the heavy strand (OH sequences) can be identified in six contigs. Multiple coding sequences for the intronic endonucleases GIY and LAGLI can be identified in six contigs. Strikingly, both between *N. caninum* strains, or between *N. caninum* and *T. gondii*, no single contig bearing the same combination or order of gene coding fragments could be identified (**Fig 5**).

**Figure 5.**
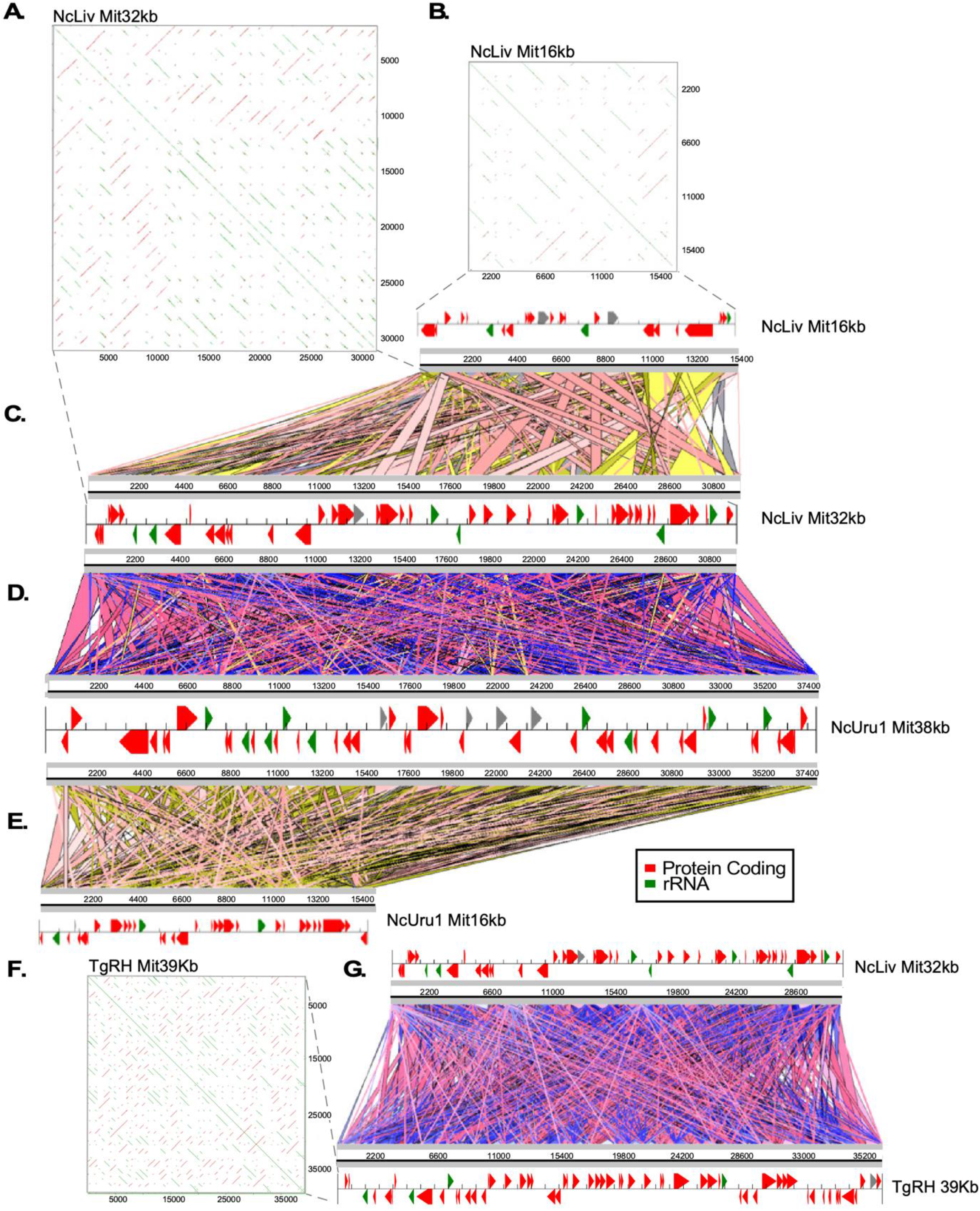
Comparative analysis of mitochondrial genome structures and annotations of *Neospora* and *Toxoplasma* reveals gene fragmentation and reshuffling between species and strains. **A.** The repetitive nature of the gene structure in a 32kb mitochondrial DNA contig of *Nc Liverpool* is graphically represented in a YASS plot. **B.** The repetitive nature of the gene structure in a 16kb mitochondrial DNA contig of *Nc Liverpool* is graphically represented in a YASS plot. **C.** Comparative alignment between two NcLiv mitochondrial contigs of 16 and 32kb, respectively. **D**. Comparative alignment between an NcLiv mitochondrial contigs of 32kb and an NcUru1 mitochondrial contig of 38kb. **E**. Comparative alignment between two NcUru1 mitochondrial contigs of 16 and 38kb, respectively. **F.** The repetitive nature of the gene structure in a 16kb mitochondrial DNA contig of *NcUru1* is graphically represented in a YASS plot. **G.** Comparative alignment between an *NcLiv* mitochondrial contigs of 32kb and an *T. gondii* mitochondrial contigs of 39kb.

## DISCUSSION

Our ability to sequence genomes and annotate genes have greatly enhanced our understanding of the molecular basis of life, health and disease. Comparative genomics has allowed us to establish evolutionary relationships among living organisms, and aided in the development of specific molecular diagnosis and the rational prediction of selective drug targets. Widely used sequencing technologies, such as Sanger, 454 and Illumina, have played a pivotal part in these advancements. However, the limitations of these technologies, namely their trouble reading through repetitive regions and their short read outputs, have led to assembly artifacts that are currently widely distributed in genome and proteome databases^43^. A number of protozoan parasite genomes have been recently revisited using third generation sequencing technologies. One noteworthy example is the genome of *Trypanosoma cruzi*, the causative agent of Chagas disease. This genome was massively improved for several strains. Its assembly went from being highly fragmented, in over 4000 short contigs, to less than 2000 contigs, doubling the overall genome size^44,45^.

Such improvements have allowed accurate account for gene copy number and the identification of an underlying genome structure. Such advances are essential in order to profiting from the advent of highly specific “guided” genome editing technologies, such as CRISPR/Cas9 technology, understand virulence linked to gene copy number, and inter and intra-species differences.

Here, we disentangled the genomes of two closely related species; *Neospora caninum* and *Toxoplasma gondii* by *de novo* assembling them. The *N. caninum* genome had been previously assembled under the presumption that they were largely syntenic, using short reads^6^. We uncovered that even though half of the genome is indeed structured quite similarly between *N. caninum* and *T. gondii,* seven out of thirteen chromosomes differ significantly from each other. Importantly, our unbiased assembly of both genomes revealed that, strikingly, the karyotype of these apicomplexans is of 13 chromosomes. Our results suggest that chromosomes previously mapped as ChrVIIb and VIII are, in fact, a single chromosome. A number of previous findings support this fusion. Several linkage analysis assays to uncover virulence factors demonstrated that these chromosomes always segregate together^46,47^. Genome interactions mapped by Hi-c analysis revealed chrVIIb and chrVIII had a higher number of interactions with each other than any other combination of chromosomes did. The latter study also showed that the number of contacts between the right telomere of chrVIIb and the left telomere of chrVIII were the highest of all, suggesting that these could be physically linked^48^. In addition, identification of the centromere of ChrVIIb was unattainable by ChIP-Chip or ChIP-seq^32,49^.

Interestingly, we detect a large inversion in ChrXII of *N. caninum* with respect to the *T. gondii* chromosome. This inversion is suggested in the 3D analysis of genome structure done by Hi-C using the *Toxoplasma gondii* Type II strain ME49^48^. However, we do not detect such inversion in our *T. gondii* RH genome assembly suggesting that the ChrXII inversion could be strain-specific.

Genome structure differences and rearrangements are widely observed amongst apicomplexans^1^. However, the driving forces of these differences are ill understood. It is well established that genomic divergence amongst Trypanosomatids, for example, can be partially ascribed to the presence of transposable elements within their genomes. Apicomplexans’ genomes, however, are devoid of such sequences.^1^ We identified low complexity, repetitive regions but their appearance did not correlate with recombination prone chromosomes, but were rather evenly distributed throughout the genome. We did, however, identify a number of repetitive motifs frequently located in the vicinity of regions where synteny is lost. Experimental validation of these motifs as drivers or “soft spots” for recombination would be needed to mechanistically link them to chromosomal rearrangements.

Unlike the situation for *T. gondii* whereby multiple strains have been fully sequenced, whole genomes of *N. caninum* were so far limited to the reference strain *N. caninum Liverpool*. Detailed population genomics studies based on whole-genome sequences from multiple strains worldwide are lacking, and so is our current understanding of population genetic structure of *Neospora.* Very recently, however, a study analyzing 19 linked and unlinked genetic markers of 50 isolates collected worldwide resolved a single genotype of *N. caninum*^50^. This is consistent with our results whereby our whole genome assembly of NcUru1 is practically indistinguishable from that of NcLiv, despite deriving from completely different geographical locations (Europe vs. South America). Nonetheless, it is well established that great genetic variability exists amongst *N. caninum* strains in the form of SNPs at particular loci and that these genetic differences underlie phenotypic variability^12,51,52^. In this context, it is noteworthy that linkage assays, whereby these minimal differences can be correlated to virulence phenotypes, rely on correct genome assemblies.

The apicoplast is a validated drug target to fight apicomplexan caused diseases such as toxoplasmosis and malaria^34,53^. Despite its importance, however, very few complete apicomplexan apicoplast genome sequences have been reported. The *N. caninum* plastid genome sequence had not been identified prior to this study. However, plastid genome physical characterization suggested a size of approximately 35 kb; whereby formation of oligomeric molecules, migrating as linear molecules in approximate multiples of the unit length, was detected^54^. Despite the ample coverage, and average read lengths that comfortably exceeding the putative apicoplast genome size, we were not able to find evidence that preferentially supported a circularize plastid genome of 35kb over two linear molecules adding up to over a 60kb, fostering an inverted repeat on their ends. It is feasible, however, that the comparatively lower GC content of the apicoplast genome hinders its effective sequencing yielding lower coverage. Nonetheless, our sequencing data cannot distinguish between the presence of a circular molecule harboring an inverted repeat or of two linear molecules, of high identity to each other. Further experiments would be required to elucidate this conundrum.

On the other hand, the apicomplexan mitochondrial genome had been described before as consisting of repeated elements of 6-7kb in length^55^. Mitochondrial genomes commonly encode a number of proteins required for its maintenance; part of the translation apparatus, tRNAs, large and small ribosomal RNAs *rns* and *rnl*; membrane associated proteins which catalyze oxidative phosphorylation: cytochrome b (*cob*), subunits of cytochrome c oxidase (*cox*), ATP synthase subunits 6, 8 and 9, subunits of the NADH dehydrogenase complex; and additional ORFs of unknown functions.

Here, we found that the mitochondrial genomes of *N. caninum* code for fragments of cob, cox1, cox3, rrns, rrnl, Lagli and Giy.

Interestingly, while the nuclear and apicoplast genomes are virtually identical between the two *N. caninum* strains sequenced, mitochondria genomes are strikingly different. Noteworthy, different combinations of ORFs are observed between species (*N. caninum* and *T. gondii*), and between strains of *N. caninum*. Mitochondria also differed greatly in size. Varying size mtDNA fragments had been inadvertently observed before for *Eimeria tenella*, whereby southern blot of Cox3 yielded a smear pattern of 4 to 20 kb^56^. In addition, despite our ample sequencing coverage, we did not find any circular molecule, suggesting that the mtDNA is composed of linear fragments. This has been reported for *Babesia*, *Theileria* and *Plasmodium* mtDNA^57,58^. Gene structure organization differences are also observable amongst these closely related hematozoa apicomplexans.

Strikingly, no mitochondrial contig in *N. caninum* nor *T. gondii* contained a fully functional copy of *cox1* or *cob*. Likewise, *Plasmodium* mitochondrial ribosomal RNAs display high degree of fragmentation. It is unclear how these small RNA fragments come together to form a ribosome^59^. A similar gene structure of fragmented gene pieces (divided in modules) has been described in the free living kinetoplastids. The mitochondrial gene modules of the unicellular flagellate *Diplonemid* are separately transcribed, followed by the joining of partial transcripts to contiguous RNAs^60^. Mitochondrial gene editing, whereby transcripts are altered post-transcriptionally to render a functional product, has so far not described for apicomplexans. However, this process is quite common and mechanistically diverse in this organelle in a wide range of species ranging from protozoa to plants. On the other hand, functional copies of mitochondrial proteins could be encoded for in the nuclear genome and imported into the organelle. Protein import is largely used by the apicoplast, whereby proteins required for its metabolic functions are all imported from the nucleus^38–41,61^. Only those proteins required for the maintenance of its genome are fully transcribed and translated within the organelle^42^. Mitochondrial tRNAs, and ATP synthase subunits are known to be imported in *T. gondii*^55,62,63^. Noteworthy, a functional *cox1* copy has been annotated in the *T. gondii* nuclear genome (TgME49_209260), and a homolog is present in the reference NcLiv genome (NCLIV_003650). In addition, nuclear-encoded divergent cox-related proteins have been identified in the mitochondrial proteome of *T. gondii*^64^.

Nonetheless, our results pose questions regarding the mechanisms of mitochondrial protein synthesis and transport which merit consideration.

The molecular mechanisms underpinning such high variability of genetic content among mitochondrial genomes are unknown. However, we identified the presence of LAGLI and GIY among the mitochondria encoded genes. Both these proteins are endonucleases encoded for in invasive introns shown to be mobile elements. Intraspecific variability of fungal mitochondrial genomes has been mechanistically linked to the movement of these endonucleases^65^. Likewise, a GIY type endonuclease present in the second intron of the mitochondrial *cytochrome b* gene in the fungus *Podospora curvicolla* was shown to autonomously transfer from an ORF-containing intron to an ORF-less allele^66^. Our results pave the way for exploring the exact contribution of these endonucleases to genome variability. It remains to be determined, as well, whether the mitochondrial genome variability observed corresponds to a clonal collection of multiple fragment, homogenously present within a population, or whether the contigs assembled represent a cohort of heteroplasmic mtDNA differentially distributed at the population level.

Overall, our results highlight distinct nuclear genomic structures, and a variable mitochondrial genome, as previously unexplored sources of genetic variability among apicomplexans. Variability is not only observed amongst closely related species, but also between strains. Efforts to explore the mechanistic contributions of this variability could shed light onto the molecular underpinnings of virulence related traits ranging from fitness to differences in drug susceptibility.

## ACKNOWLEDGEMENTS

We thank Dr. Fernando Alvarez and Dr. Jon Boyle for critical reading of this manuscript. This work was funded by a grant from the National Agency for Research and Innovation (ANII) and the National Institute for Agricultural Research (INIA) number FSSA_X_2014_1_106026 awarded to CR, and an installment grant from the Banco de Seguros del Estado (BSE) awarded to MEF.

## AUTHOR DECLARATION

All authors have read and approved the current version of this manuscript and have agreed to its submission

